# Halophilic nematodes live in America’s Dead Sea

**DOI:** 10.1101/2023.04.12.536621

**Authors:** Julie Jung, Tobias Loschko, Shelley Reich, Michael S. Werner

## Abstract

Extremophiles can reveal the origins of life on Earth and the possibility of life elsewhere. Most identified extremophiles are single-cell microbes, leaving gaps in our knowledge concerning the origins and habitable limits of multicellular organisms. Here, we report the recovery of roundworms (Phylum Nematoda) from the Great Salt Lake (GSL), UT, a hypersaline lake referred to as “America’s Dead Sea”. Nematodes were found primarily in microbialites, benthic organosedimentary structures once abundant on early Earth. 16S sequencing of individual nematodes revealed a diverse bacterial community distinct from its surrounding habitat. Phylogenetic divergence compared to Owens Lake, another terminal lake in the Great Basin, suggests that GSL nematodes represent multiple previously undescribed species. These findings update our understanding of halophile ecosystems and the habitable limit of animals.

**One-Sentence Summary:** We report the discovery of novel halophilic nematodes in microbialites of the Great Salt Lake, UT.

## Main Text

The Great Salt Lake (GSL) in northeast Utah fluctuates around ∼15% salinity in its southern arm and 30% in its northern arm, making it one the most saline bodies of water on Earth. Early non-indigenous settlers assumed it was too saline to harbor life (*1*), but modern methods have revealed a thriving extremophile community inhabiting the lake’s benthic zone (*2–4*). The GSL’s shallow marginal areas are home to salt-tolerant microorganisms that build calcium carbonate mounds up to ∼ 1 m in diameter called microbialites (*2, 3, 5, 6*). These reef-like structures span 700-1000 km^2^, or 20% of the lake bottom, making it one of the largest assemblages of microbialites in the world. While rare today, microbialites were the dominant macroscopic evidence of life on Earth for two billion years (*7, 8*). Understanding the formation and biota of these structures could provide important clues to the origin and ecology of early life on our planet.

Despite the lake’s flourishing microbial community, only two animal taxa have persisted in its benthic zone: brine shrimp (*Artemia sp*.) and brine fly larvae (*Ephydra sp*.). These invertebrates, which feed on microbialite bacteria, are a critical food source for the millions of migratory birds passing through the Pacific and Central flyways annually (*2, 9, 10*). Yet, like many of the world’s saline lakes, the GSL is shrinking at an alarming rate (*11, 12*). Decades of water diversion exacerbated by severe drought in the region have caused historic lows in water elevation (*13, 14*). As a result, exposed microbialite habitat and record-high salinity levels threaten both benthic zone inhabitants and the upper trophic levels that depend on them. Thus, there is a pressing need to understand this lynchpin community and the limits of their habitability.

Although nematodes had never been described in the GSL, several lines of evidence motivated further investigation. As the most abundant animal phylum on the ocean floor and terrestrial biosphere, nematodes exhibit remarkable diversity with at least 250,000 extant species (*15–17*). Some of these have been found kilometers below the surface (*18*), and in extreme cold and arid conditions (*19*). Moreover, a recent study reported several nematode species in Mono Lake, a saline and arsenic-rich terminal lake in California (*20, 21*).

Here, we report the recovery of nematodes from the benthic zone of the GSL – representing the most saline environment nematodes have ever been recovered from – and describe their preferred habitat, microbiome, and phylogenetic relationships. These findings raise the number of metazoan taxa in the GSL benthos from two to three, extend the known habitable zone of nematodes, and provide a foundation for future studies of extremophile animal–microbe communities.

### Halophilic nematodes live in the Great Salt Lake

We investigated whether nematodes live in the GSL by conducting seasonal sampling from the Fall of 2021 to the Summer of 2022. Our field sites spanned a salinity gradient that started in the freshwater Weber River (sites 1 and 2), transitioned to a brackish delta (site 3), and terminated in three sites within the south arm of the GSL (sites 4–6; Fig. 1A, S1).

**Fig. 1.**
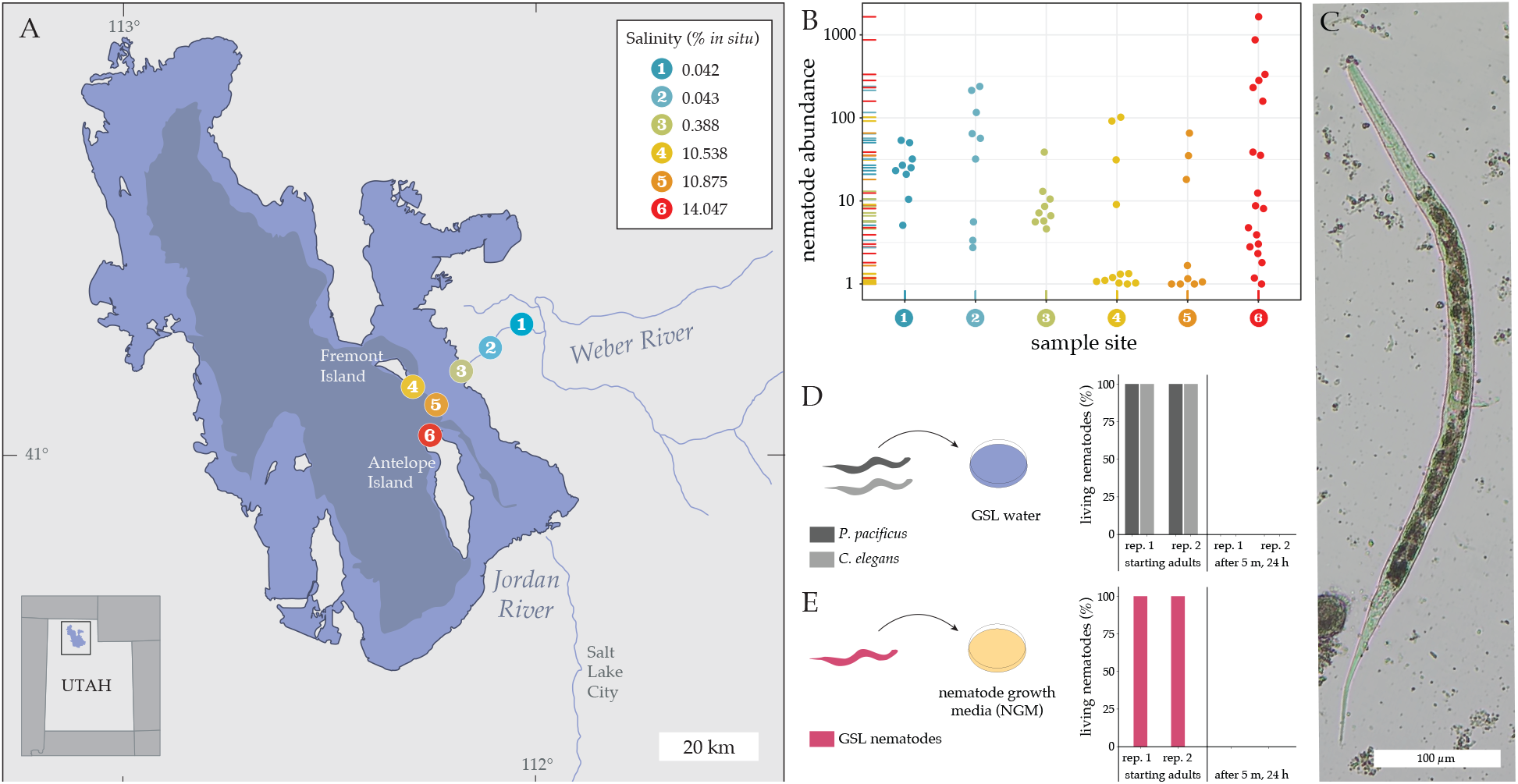
Nematodes live in the Great Salt Lake, Utah. **(A)** Locations and average *in situ* salinity of the six sampling sites around (1-3) and within (4-6) the Great Salt Lake. Lake topographies depict NASA SRTM2 v.2 data from 2007 and UGRC LiDAR data from 2016. Samples were collected in spring, summer, and fall of 2021 and the summer of 2022. **(B)** Nematode abundance measured in number of nematodes per 100g of dry lake sediment on a log scale and colored by sample site from which samples were collected, spread horizontally to show stacked points. **(C)** Representative nematode extracted from the south arm of the Great Salt Lake on March 3, 2021. **(D)** Survival of *C. elegans* and *P. pacificus* before and after being placed in Great Salt Lake water for 5 minutes vs. **(E)** survival of nematodes extracted from GSL before and after being placed on NGM plates for 5 minutes.

We were unable to recover nematodes in the GSL using the standard Baermann funnel technique over several sampling trips. However, when we used a sucrose-density centrifugation method, which can process larger samples and has historically been used to isolate nematodes from low density sites in desert and polar regions (*19*), we consistently recovered live nematodes from every site (Fig. 1B-C, Movie S1). To confirm whether GSL nematodes were true halophiles, we compared their viability in GSL water to that of two common laboratory nematodes: *Caenorhabditis elegans* and *Pristionchus pacificus*. Most GSL nematodes were alive and motile for over a week in the water from which they were recovered. In contrast, all *C. elegans* and *P. pacificus* died within five minutes of being placed in GSL water (Fig. 1D). Importantly, the converse was also true; GSL nematodes died within five minutes when placed on standard nematode growth media plates (Fig. 1E), indicating specific adaptation to the environment of the GSL.

Average nematode abundance in the GSL was lower than typically found in terrestrial ecosystems, but similar to the densities found in Mono Lake (Table S1) (*17, 20*). Surprisingly, nematode abundances in hypersaline GSL sites were comparable to densities in nearby freshwater Weber River sites (f=3.05, p=0.09). Consistent with this observation, salinity did not drive differences in nematode abundance across sample sites (f=0.028, p=0.879, Fig. 2A). Salinity above 15% has been shown to negatively affect the microbial mat community and brine shrimp and brine fly growth (*22–25*). Yet, we found stable nematode abundances in samples of up to 19% salinity. Population sizes were also stable throughout 2021 (f=1.105, p=0.338, Fig. 2B), which suggests these animals can tolerate large shifts in UV, temperature, and salinity that accompany the seasonal changes in high desert environments (*26, 27*). Nevertheless, sampling in the summer of 2022 yielded significantly fewer nematodes concurrent with a record low lake elevation and mean annual loss of 35.76 cm (Wilcoxon rank sum test, Z=2.47, p=0.00675). Thus, while these newly discovered inhabitants of the GSL benthos are remarkably resilient, the continued recession of the lake likely poses a threat to their survival.

**Fig. 2.**
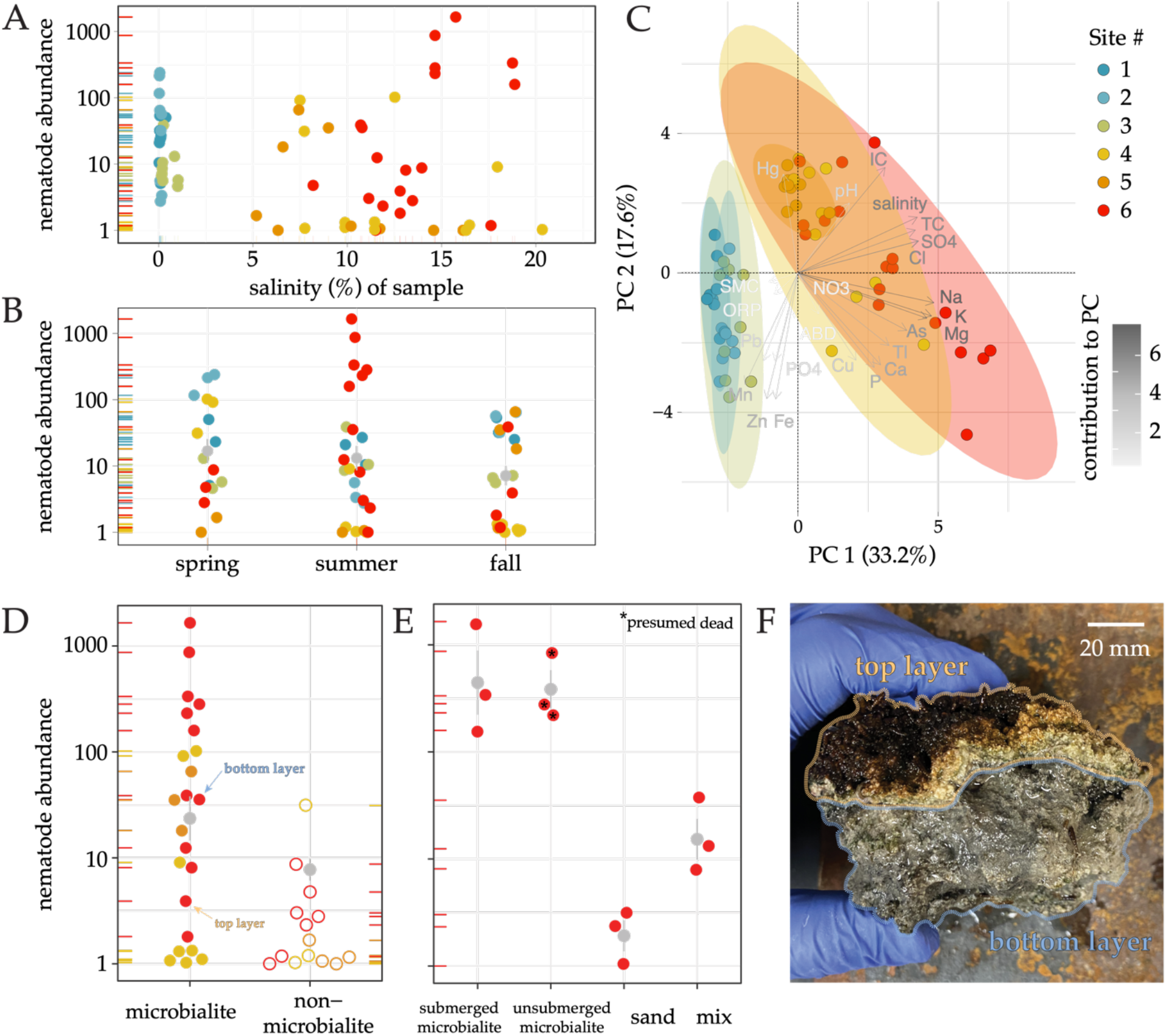
Microbialites are a microhabitat for Great Salt Lake Nematodes. Nematode abundance by **(A)** salinity of each sample and **(B)** season during which each sample was collected. **(C)** Principal component analysis of samples by site. Color and length of line in biplots are determined by contribution to the PC. Ellipses show 95% confidence intervals. **(D)** Nematodes are preferentially found in microbialite (filled circles) compared to non-microbialite (unfilled circles) samples from GSL. We extracted nematodes from both the top layer and bottom layer of one sample, as annotated. **(E)** Nematode abundance, as collected from submerged microbialite, recently unsubmerged microbialite, submerged sand directly adjacent to microbialites, and a microbialite/sand mix. Those marked with black stars indicate all extracted nematodes were dead upon extraction. **(F)** Representative microbialite sample showing distinct top and bottom layers. Abundance was measured in number of nematodes per 100g of dry lake sediment on a log scale. Data points for categorical variables are individual samples colored by sample site from which samples were collected and spread horizontally to show stacked points. Gray summary stats for categorical variables show means and standard error.

### Microbialites are a microhabitat for GSL nematodes

In the absence of a correlation with salinity, we searched for other covariates of nematode habitability. We measured pH, soil-moisture content, oxidation-reduction potential, soluble anions and extractable elements (Na, Mg, K, Ca, P, Mn, Fe, Cu, Zn, As, Hg, Tl and Pb), and Total Organic Carbon and Inorganic Carbon (see supplementary materials and methods). Sites within the GSL had a distinct chemical profile as compared to our freshwater samples (Fig. 2C). Additionally, GSL samples showed concerning trends of mercury and high levels of arsenic and thallium, which represent well-known health hazards if the lakebed becomes exposed as they can be distributed by wind and atmospheric deposition to the surrounding metropolitan area (*28, 29*). A thorough analysis of these and other environmental variables will be presented in a compendium manuscript. However, regarding our original question, we were unable to detect a significant correlation between any chemical variable and nematode abundance (Fig. S2, Table S2).

We expanded our search for a signature of nematode habitability by examining potential differences in microhabitat. Through our sampling efforts, we noticed higher numbers of nematodes from microbialites (Wilcoxon rank sum test, Z=3.55, p=0.0002, Fig. 2D). To further investigate this association, we homogenized microbialite samples and passed the crushed sediment through size-selective sieves. Crushed microbialite sediment yielded on average 237x more living nematodes than did sediment directly adjacent to microbialite mounds, while a mix of microbialite and non-microbialite sediment exhibited intermediate nematode abundances (Fig. 2E). Nematodes were found in equally high abundances from both submerged and recently (within 12 days) unsubmerged microbialites, but all nematodes extracted from dry microbialites were immobile, rigid, and presumed dead (Fig. 2E).

The microbialites sampled in this study subscribed to previously described morphology, each showing a succession of thin orange and porous green layers (Fig. 2F) (*22*). The top layer exhibited clusters of photosynthetically active coccoid cyanobacteria and carotenoids. This upper mat quenches UV rays and protects the deeper microbialite layers (*6*). Intriguingly, 2.7-fold more nematodes were found in the bottom layer than the upper mat (Fig. 2D). Taken together, GSL nematodes are preferentially found in active microbialites, which may provide protection from radiation and desiccation.

### The microbiome of Great Salt Lake nematodes hints at co-evolutionary mechanisms of survival in hypersaline and arsenic-heavy environments

Nematodes may also feed on the bacteria that are present in microbialites. Indeed, the nematodes identified from our sampling were likely bacterivores based on their relatively narrow and smoothly lined buccal cavities (Fig. S3) (*30*). Endo- and ecto-symbiotic (i.e., cuticular) bacteria could also provide niche-specific functions for their nematode hosts as in other extreme environments (*31, 32*). To identify bacterial food sources in microbialites, and potential symbiotic relationships, we sequenced the V4 region of bacterial 16S rRNA using individual nematodes as input. The median Shannon alpha diversity of bacteria across 15 nematodes was 4.8 and the median diversity among archaea was 2.2 (Fig. 3A, S4), demonstrating a relatively rich microbial community, especially compared to other hypersaline lakes (*33*). Importantly, rarefaction curves indicated that our analysis captured all the diversity of the true microbial community (Fig. S5). Moreover, our ‘no nematode’ control samples did not exhibit PCR amplification, arguing that the associated microbial taxa resided either on or inside the worms (Fig. S6).

**Fig. 3.**
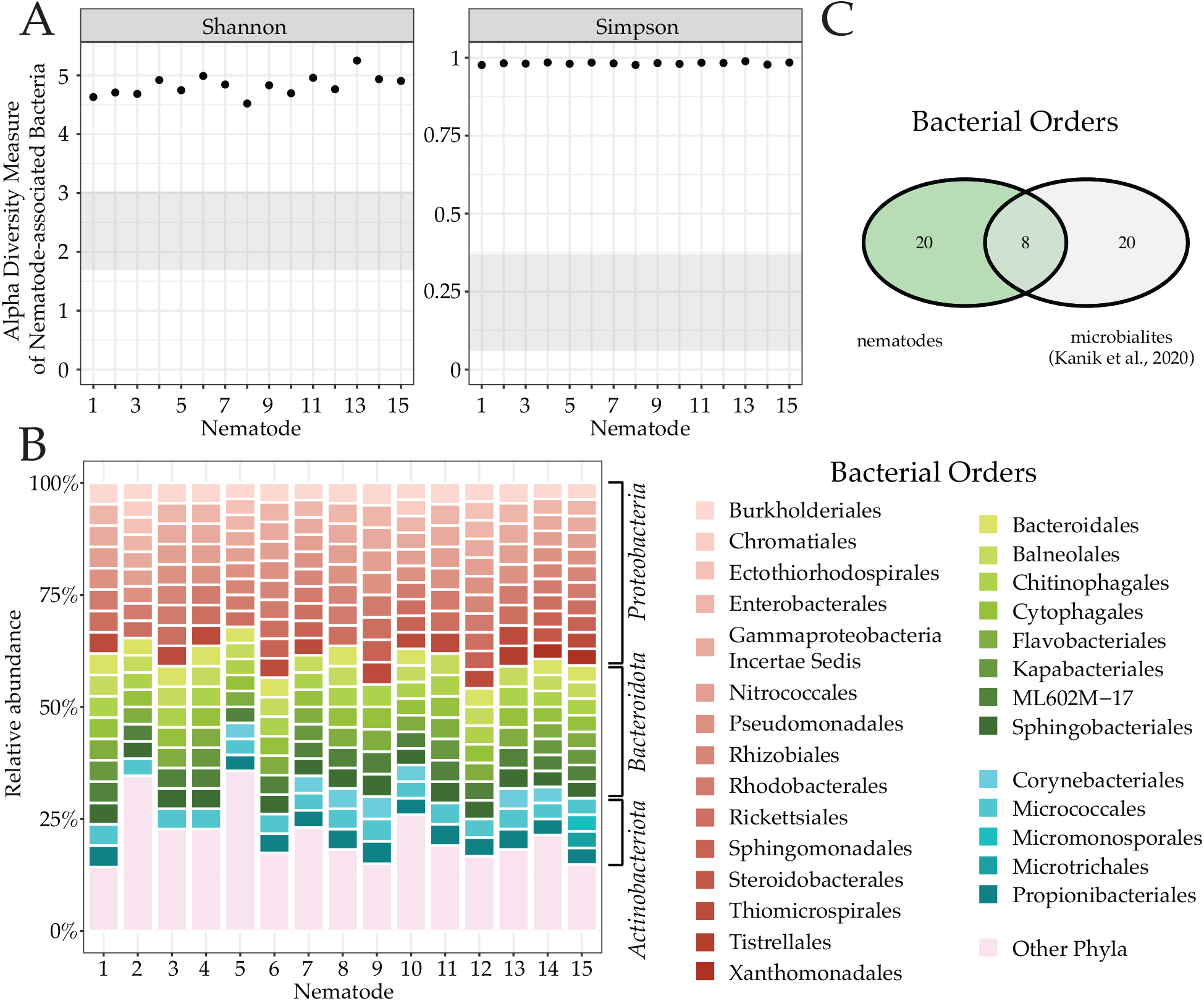
Microbiome of Great Salt Lake nematodes. **(A)** Shannon and Simpson diversity indices of nematode-associated bacteria. Each dot represents an individual nematode from which associates were analyzed. Gray box shows range of diversity indices in hypersaline lakes in Western Mongolia (*33*). **(B)** Overview of nematode-associated microbes. The relative abundance of nematode-associated bacterial communities binned by order. All 15 nematodes were collected from Fremont Island (site 4) and taxonomic bins are grouped by phylum. **(C)** Venn diagram of bacterial orders specific to Great Salt Lake nematodes, specific to Great Salt Lake microbialites (*3*), and shared between the two.

Photosynthetic bacteria are typically dominant in the surface layers of microbialites whereas chemo-heterotrophs and anaerobes are dominant in the deeper layers (*6, 34, 35*). Consistent with greater nematode abundance in the deeper layer of microbialites, only 4% of amplicon sequence variants (ASVs) mapped to cyanobacteria. Several of the remaining bacterial orders overlapped with orders found in GSL microbialites (Fig. 3B) (*3*). For example, Proteobacteria were one of the dominant phyla found to be associated with GSL nematodes and are known to deposit carbonate by sulfate reduction for microbialite formation (*26*). Strikingly however, ∼70% (20/28) of bacterial orders were specific to nematodes and not found in microbialites (Fig. 3C). Thus, single-worm 16S-sequencing revealed a rich and largely distinct nematode-microbial community.

The dominant taxa associated with GSL nematodes were *Psychroflexus* and *Candidatus Aquiluna*, which belong to the most abundant bacterial Orders Flavobacteriales and Micrococcales, respectively (Fig. S7A). These are known to act as secondary producers by supplementing their energy metabolism using light-harvesting rhodopsin pigments and other photosynthetic reaction centers (*3, 26, 36*). GSL nematodes also associated with the halotolerant bacterium *Halomonas elongata* (Fig. S7A) and several halophilic archaea known to regulate the amount of salt and osmotic pressure inside their cells by keeping charged solutes and amino acids on their cell surfaces (Fig. S7B) (*37–40*). Members of the dominant archaeal phylum present in GSL nematodes, Halobacterota, are known to use arsenic compounds for bioenergetic processes (*41, 42*) and have adopted a variety of resistance mechanisms to overcome arsenic stress (Fig. S7B) (*43*). More than a third of GSL samples exceeded the threshold effect concentration of arsenic (10 μg/g), and the highest exceeded the limit by 1100% (Fig. S2L), which likely yields extensive ecological effects at multiple trophic levels (*44*). Samples collected from site 6 were particularly high in arsenic content, which can be attributed to that site’s relative proximity to mining and smelting activities (*45*). Taken together, the GSL-nematode microbiome contains several members with known mechanisms of energy production, hypersaline tolerance, and toxic-chemical resistance. These associations may reflect ancient animal–microbe interactions within microbialites, and are attractive candidates to investigate habitat-adapted symbiosis or co-evolution to the extreme environment of the GSL.

### Genetic diversity of Great Salt Lake nematodes

Next, we performed small subunit ribosomal RNA (SSU) sequencing to genotype nematodes from each site (Table S3) (*46*). Nematodes found in our freshwater and mostly freshwater sites (1-3) belonged to 18 different families. However, all but two nematodes extracted from the south arm of the GSL belonged to a single family: Monhysteridae (Fig. 4). With more than 200 species and global distribution across marine and estuary sediment, Monhysteridae are likely an ancient nematode family (*47, 48*), categorized as microbivorous for purposes of functional and trophic ecology (*49*). Monhysterids are also one of the most abundant nematode families in deep-sea hydrothermal vents and are the dominant metazoan in abyssal zone sediment (*50–52*), demonstrating an ability to adapt to extreme environments. The dramatic drop in diversity in sites 4-6 corresponds with the transition from brackish delta to hypersaline sites in the lake, indicating adaptation of specific lineages to the extreme salinity.

**Fig. 4.**
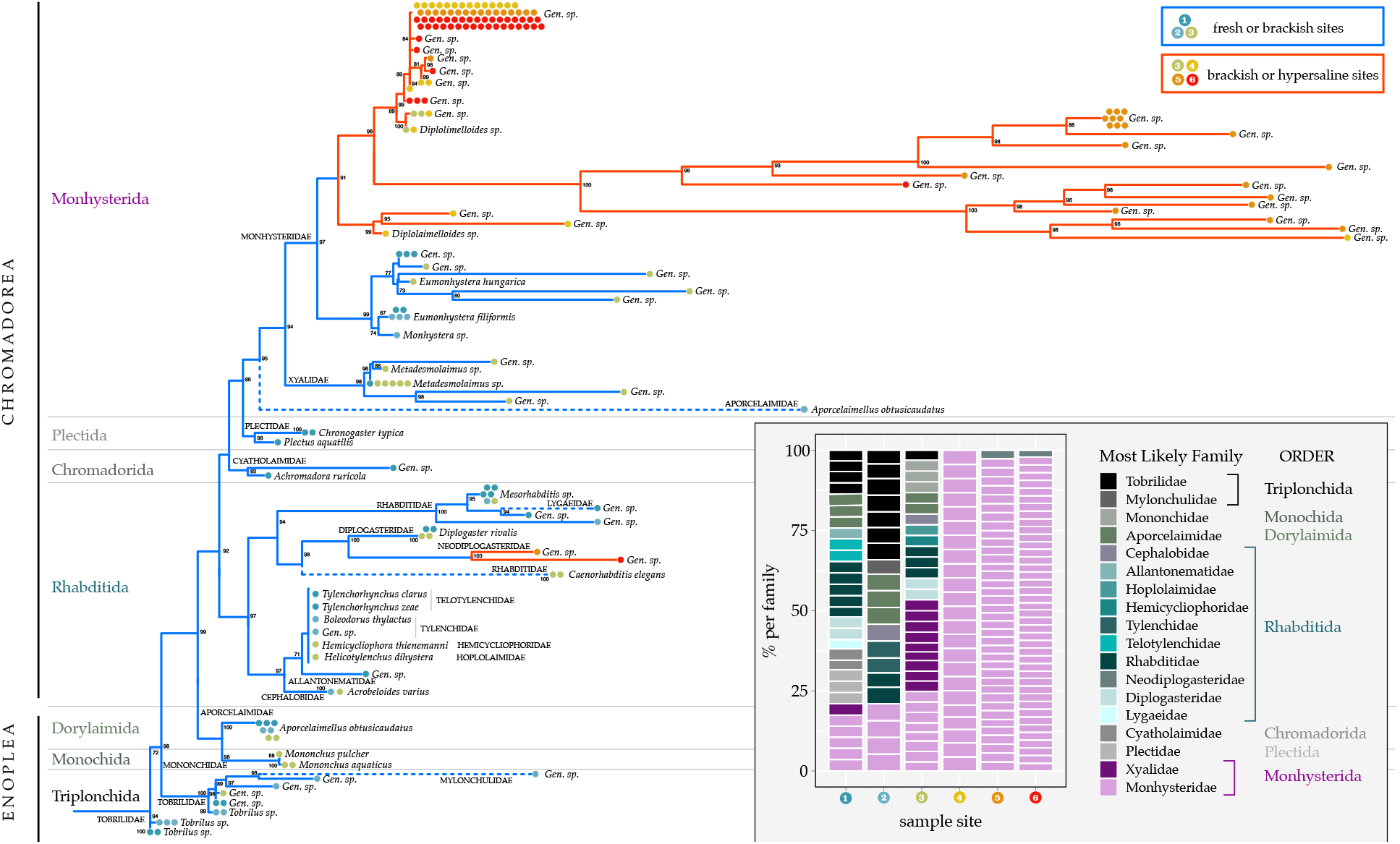
Diversity of Great Salt Lake nematodes. Maximum likelihood phylogenetic tree of isolated nematodes, based on aligned SSU sequences and a GTR model for nucleotide substitutions. Tree tips denote the best SSU BLAST hit (NCBI) >97% identity. Nodal support for analysis (shown in black) was performed for species identification with 1,000 bootstrap replicates. Nodes with support values below 0.70 within a single family were collapsed into polytomies. Clades consisting of multiple individuals of a single SSU haplotype were collapsed into a single branch and labelled with sample sizes (each point is 1 nematode) and which site that nematode was collected from. (**Inset)** Nematode family composition by site. Each box is an individual nematode, colored by the most likely family and order.

To investigate the evolutionary history of GSL nematodes we constructed a Maximum Likelihood phylogenetic tree based on single-worm SSU sequences. Our tree recapitulated established phylogeny (*46, 53*), and included two of the three major lineages that exist within phylum Nematoda, class Enoplea and class Chromadorea (Fig. 4). Importantly, a tree made from a second 18S locus exhibited the same overall topology (Fig. S8). Thus, while analyzing more loci is needed to make fine-scale evolutionary inferences, we felt confident that these data could reveal the general phylogenetic relationship of GSL nematodes.

We observed at least two well-defined clades within Monhysteridae. All 103 nematodes in the first clade were collected from hypersaline to brackish sites (sites 3, 4, 5, and 6), while all 14 nematodes in the second clade were collected from freshwater to brackish sites (sites 1, 2, and 3) (Fig. 4). Monhysterids from the GSL also differed morphologically from Monhysteridae worms from our freshwater samples (Fig. S3). Freshwater Monhysteridae were larger (Fig. S3A) and displayed two conspicuous cephalic setae (Fig. S3B-C), which were missing or substantially smaller in GSL Monhysterids (Fig. S3D-F) (*30*). Two specimens in the hypersaline clades appeared to be from the genus *Diplolaimella*, a free-living group that typically inhabits marine and coastal sediment. However, the vast majority of nematodes from sites 4-6 exhibited <97% best-hit match on NCBI. Morphological comparisons are ongoing to determine their precise alpha taxonomy. Yet, the combination of (1) adaptation to extreme salinity, (2) large sequence divergence and (3) evidence of multiple branches suggests a prolonged period of reproductive isolation. We interpret these data as at least two, if not several, novel species of nematodes unique to the south arm of the GSL.

### Comparison to an analogous terminal saline lake in the Great Basin

To compare nematode abundances and investigate the potential evolutionary history of GSL nematodes, we collected submerged benthic sediment from six sites in Owens Lake, a terminal lake on the western edge of the Great Basin (Fig. 5A). Once one of the largest inland bodies of water in the US, Owens Lake was drained by 1926 to accommodate the burgeoning water demand of Los Angeles. As a result, dust storms from winds blowing off the Sierra Nevada mountains swept up toxic particulates from the dry lakebed, creating the nation’s worst source of dust pollution. In 2006, the Los Angeles Department of Water & Power began an expansive effort to mitigate this crisis, ranging from gravel cover and tilled soil to managed vegetation and shallow flooding (Fig. 5A). We recovered 15-257 nematodes per 100g of sediment from the three least saline sites, where measured salinity was less than 1% (Fig. 5A-B). However, the remaining sites – which ranged from 4.3% to 13.4% salinity – yielded nearly negligible nematode abundances (Fig. 5B). Thus, in contrast to the GSL, nematode abundance in Owens Lake was inversely correlated with salinity (Fig. 5B).

**Fig. 5.**
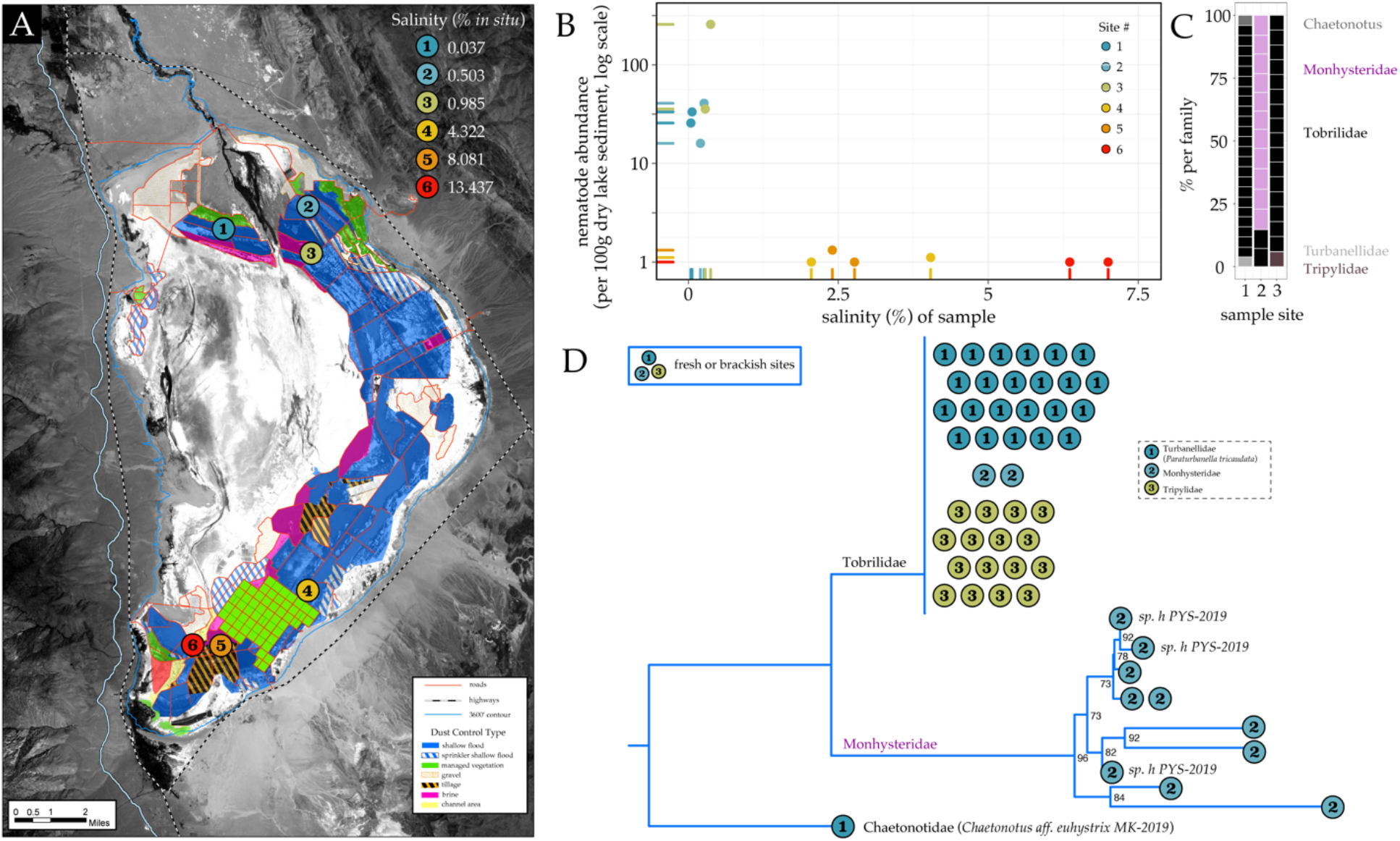
Comparison of nematodes isolated from Owens Lake, CA. **(A)** Locations and *in-situ* salinities of the six sampling sites within the restored Owens Lake. Original map credit: LADWP. **(B)** Nematode abundance by salinity of each sample. **(C)** Nematode family composition by site. Each box is an individual nematode, colored by the most likely family. **(D)** Maximum likelihood phylogenetic tree of the isolated nematodes, based on SSU sequences and a GTR model for nucleotide substitutions. Tree tips denote the best SSU BLAST hit (NCBI) >97% identity. Nodal support for analysis (shown in black) was performed for species identification with 1,000 bootstrap replicates. Nodes with support values below 0.70 within a single family were collapsed into polytomies. Clades consisting of multiple individuals of a single species were collapsed into a single branch and labelled with sample sizes (each point is 1 nematode) and which site that nematode was collected from.

We also discovered notably less diversity at the freshwater sites within Owens Lake compared to freshwater input sites near the GSL. Nematodes from sites 1 and 3 were primarily in the family Tobrillidae, while nematodes from site 2 were primarily in the family Monhysteridae (Fig. 5C). Interestingly, the Monhysterids found in site 2 seem to be the same SSU haplotype (sp. h PYS-2019) discovered in 2016 in Mono Lake, located just 140 miles northwest of our sites (*20*). Given the similar geology between the GSL and Owens Lake, the large differences in nematode abundance and taxa were notable. We attribute these differences to two non-mutually exclusive possibilities: decades of desiccation caused a major local extinction event in Lake Owens, or the microbialite mounds within the GSL provided a protective habitat for nematodes to thrive in hypersaline environments. In either case, sampling data from Lake Owens suggest that the continued survival of GSL nematodes depends on the water level of the GSL.

## Conclusion

The GSL is an extreme hypersaline environment with only two currently known metazoan groups, brine flies and brine shrimp. Here, we report the discovery of halophilic nematodes in the benthic zone of the south arm (10-20% salinity), representing the most saline environment from which nematodes have been recovered. Phylogenetic analysis from 18S sequencing data of recovered nematodes suggests that they represent multiple previously undescribed species. Collectively, our findings expand the biodiversity of the GSL and introduce a new system to study the ecology and evolution of extremophile animals in early-Earth analog environments.

The major habitat of GSL nematodes are microbialites – a critical source of primary production for the entire GSL ecosystem (*2*). Nematodes likely benefit from grazing on the bacteria in microbialites and may also gain protection from UV exposure or dehydration. It’s currently unclear whether the relationship is reciprocally beneficial towards microbialites and their microbial communities, but nematodes are known to contribute to niche construction through grazing, nutrient cycling, and bioturbation in diverse environments (*15, 54*). The microbiome of nematodes has significant overlap with bacteria in microbialites, and going forward, a particular focus will be applied to whether the breakdown of certain microbes by nematodes facilitates the accumulation of sediment layers. Notably though, the majority of identified microbes were distinct to the nematode host. Future studies will also investigate whether these relationships have facilitated survival in the extreme environment of the GSL.

The discovery of nematodes in the GSL presents several evolutionary questions, especially given historical fluctuations in the lake’s size and salinity levels. For much of the last 800,000 years, our sampling sites have likely stood in a saline lake (*55*). However, approximately 30,000 years ago the GSL was part of a vast freshwater lake that covered most of western Utah named Lake Bonneville. The end of the last ice-age brought a warmer and drier climate, which in addition to a natural dam break leading to a multi-year flood, formed the boundaries of the modern GSL and returned the lake to saline conditions (*56*). The American West is also currently experiencing a ‘megadrought’ considered to be the worst long-term regional drought in the past 1,200 years (*13, 14*). The drought has been exacerbated by human water consumption, driving lake levels to historic lows and salinity to historic highs. Thus, the long-term inhabitants of the GSL have had to face severe evolutionary pressures from repeated environmental change. Understanding the establishment and evolution of hypersaline benthic ecosystems – including colonization history and co-evolution with microbialites – could help us trace the boundaries of life on our changing planet, with implications for where complex life may be found elsewhere.

## Acknowledgments

We thank Byron Adams for demonstrating sucrose-density centrifugation and follow-up discussions. We also thank Talia Karasov and Effie Symeonidi for discussions of 16S sequencing, Rodolfo Probst and Jack Longino for discussions of 18S phylogeny, Sara Weinstein for discussions of 18S primer troubleshooting, William Brazelton for use of his lab’s conductivity meter, Diego Fernandez for assistance with environmental chemical analysis of soils, Carie Frantz and Ralf Sommer for feedback on the manuscript, and members of the Werner lab past and present (Thomas King, Audrey Brown, Madelyn Purnell, Bryce Bosworth, Collin Caldwell, Samantha Nestel, Hiraya Natividad II, Hannah Lee, Hephzibah Kaleem, and Cody Fitzgerald) for discussions of this research and/or assistance collecting samples and extracting worms. This research was conducted on the traditional and ancestral homeland of the Shoshone, Paiute, Goshute, and Ute Tribes. We affirm and support the University of Utah’s partnership with Native Nations and Urban Indian communities.

## Funding

National Science Foundation Postdoctoral Research Fellowships in Biology Program under Grant No. IOS-1354072 (JJ)

Start-up funds at University of Utah (MSW)

## Author contributions

Conceptualization: JJ, MSW

Methodology: JJ, TL, SR, MSW

Investigation: JJ, TL, SR, MSW

Visualization: JJ

Funding acquisition: JJ, MSW

Project administration: MSW

Supervision: JJ, MSW

Writing – original draft: JJ

Writing – review & editing: JJ, MSW

## Competing interests

Authors declare that they have no competing interests.

## Data and materials availability

Raw reads for 16S amplicon data have been deposited in the NCBI Sequence Read Archive (SRA) under BioProject number PRJNA927656. All data, including 18S sequences, and code to recreate analyses are publicly available through Dryad (doi:10.5061/dryad.nzs7h44wg).

## Notes

https://doi.org/10.5061/dryad.nzs7h44wg

### Competing Interest Statement

The authors have declared no competing interest.

