## Supplementary Material for "Halophilic nematodes live in America’s Dead Sea"

**This file includes:**

Materials and Methods
Figs. S1 to S8
Tables S1 to S3

### Materials and Methods

Sites and sampling

Sediment samples were collected from six sites of varying salinities in and around the Great Salt Lake (GSL, Fig. 1A, Fig. S1) in spring (May), summer (late July/early August), and fall (late October) of 2021 and summer (June to early July) of 2022. The median temperatures in the lake were 18°C in spring, 28°C in summer, and 8°C in fall. On July 26, 2021, the southern arm of the GSL hit a historic low (4191.28 feet above sea level), surpassing the previous record set in 1963. We collected samples from site 6 four days prior to and eight days after that day. On July 22, we collected three samples, each a mix of submerged microbialite and non-microbialite soil from the edge of the receding GSL at Bridger Bay. Twelve days later, on August 3, we returned to the same coordinates to collect from the same microbialites, now recently unsubmerged. At the same time, we collected from still submerged microbialites located 10-20 meters down the beach and the submerged sand from the space in between microbialite mounds. In the summer of 2022, we collected only from site 6 and along the water's receding edge.

Site 1 (41°13'2.0"N, 112°09'38.8"W) and site 2 (41°13'18.5"N, 112°12'03.4"W) are freshwater sites along the Weber River, one of the rivers flowing into the GSL. Site 3 (41°10'08.4"N, 112°11'27.5"W) is a transition zone with intermediate salinity very near the Weber River input. Site 4 (41°07'51.0"N, 112°18'21.0"W) is at the southern tip of Fremont Island within the GSL, while site 5 (41°07'3"N, 112°16'10"W) is a coastal area at the edge of the receding lake, and site 6 (41°03'18"N, 112°15'18"W) lies along the beach at Bridger Bay on the northwestern end of Antelope Island. At each site (within 50 m of the GPS point), at least three plastic buckets were filled with 400-600 mL of under-water sediment (<10 cm) using a shovel or PVC pipe cupped at one end by hand. Since the lake is too saline to use a motor, we were limited by how far we could paddle, bike, or hike in a single day. Sea kayaks were used as transport between islands in the spring and fall to sample from positions not reachable on foot or by bike. Sediment samples were returned to the lab and processed over the following week.

We also collected two non-microbialite soil samples from six sites of varying salinity within the restored Owens Lake on June 16, 2022 (Fig. 5A). Site 1 (36°30'32"N, 117°58'35"W) was freshwater at 0.037% salinity, while site 2 (36°20'23"N, 117°59'26"W) and site 3 (36°31'6"N, 117°56'16"W) were hyposaline at 0.503% and 0.985% salinity, respectively. Site 4 (36°21'54.7"N, 117°56'5.9"W) was mesosaline at 4.322% salinity, while site 5 (36°29'51"N, 117°56'3"W) and site 6 (36°20'31"N, 117°58'41"W) were hypersaline at 8.081% and 13.437% salinity, respectively. Samples were collected on foot and processed over the following week. All proper permits were acquired from the State of Utah and the Los Angeles Department of Water and Power.

Nematode isolation

Sediment samples were processed for nematodes by sucrose density centrifugation. Specifically, 100 or 200 ml of sediment was allocated to a large plastic beaker and filled to 800 ml with water from its collection site. Samples were then mixed to homogeneity with a spatula, and decanted over a 425 μM filter (Thomas Scientific, No. 40) to capture large debris, stacked on top of a 38 μM filter (Thomas Scientific, No. 400) to capture nematodes. Remaining sediment and nematodes on the 38 μM filter were backwashed using site-matched water into 50 ml conical tubes using a plastic funnel and centrifuged 500 x g for 5 minutes. Supernatant was removed with a pipette until 5 ml remained at the interphase with pelleted soil. The soil and interphase water were resuspended in ice-cold sugar water (454 g sucrose/L Deionized water) to 50 ml and mixed to homogeneity. If the pelleted soil was > 35 ml, it was aliquoted to two or more 50 ml conical tubes after resuspension, and each tube was filled with more sugar water to 50 ml. Resuspended samples were centrifuged 500 x g, 1 minute to separate sediment and invertebrates. Supernatants containing invertebrates were decanted over the 38 μM filter and washed gently with site-matched water. Finally, invertebrates were back washed through a funnel into a new 50 ml conical tube at about 15 ml total volume, which was decanted into 6 or 10 cm plates that had grid lines drawn on the bottom to facilitate counting. Nematodes and other invertebrates were visually counted using a Zeiss Stemi 508 dissecting microscope, and data were recorded in a Microsoft Excel Spreadsheet for subsequent analysis.

Soil measurements

10 ml of sediment was diluted with DI Water 1:10 (100 ml final) and mixed with a spatula to homogeneity. Dilute samples were probed with a YSI Professional Plus probe fitted with a conductivity meter to measure salinity (ppt) and Oxidative-Reductive Potential (ORP). pH was also measured on the diluted sample using an Accument AE150 probe (Fisher Scientific). For measuring dry weight, 10 ml of sediment was weighed in an autoclaved glass beaker and dried for 48 hours at >100° C, then re-weighed. Soil moisture content was calculated as: (wet weight - dry weight)/dry weight. Abundance, or number of worms per 100g of dry soil, was calculated as: (worms counted/volume of sediment sample)*(volume of sediment weighed/wet weight)*(wet weight/dry weight) *100. Approximately 1 g of dried sediment was analyzed for environmental chemical analysis using inductively coupled plasma mass spectrometry (ICP-MS), ion chromatography mass spectrometry (IC-MS), and IC. This analytical technique can be used to measure elements at trace levels in biological samples.

Determination of soluble anions

To determine the concentration of soluble anions $Cl^-$ , $NO_3^-$, $PO_4^{3-}$ and $SO_4^{2-}$, dry sediments were first leached with water such that ~250 mg of sediments and ~5 g Type I water (Element, Millipore, Burlington, MA, US) were mixed thoroughly with a vortex, disaggregated in an ultrasonic bath for 3 minutes, and centrifuged at 5000 rpm for 10 min. Next, a 0.500 mL aliquot of the supernatant was diluted with Type I water and mixed. This solution was run in an ion chromatograph (883 Metrohm, Herisau, Switzerland) alongside a calibration curve prepared from a multi-anion standard (Inorganic Ventures, Christiansburg, VA, USA). Anion standard (Fluka XXX) was used as a check standard measured alongside samples and in agreement within 5% of certified values. Solution and dilution factors were used to calculate the concentration reported for each anion, in units of milligrams of anion per kg of sediment.

Determination of extractable elements at pH = 7

Dry sediments were leached with 1 M $NH_4CH_3CO_2$ (ammonium acetate) buffer at pH = 7 for the determination of Na, Mg, K, Ca, P, Mn, Fe, Cu, Zn, As, Hg, Tl and Pb. About 100 mg of sediments were mixed with ~5 g buffer using a vortex and left for 24 h with occasional stirring.

After centrifuging at 5000 rpm for 10 min, a 0.100 mL aliquot of the supernatant was diluted with 2.4 % $HNO_3$ to 10.0 mL; 10 ppb In were added as internal standard and mixed in a polystyrene tube. Determination of Na, Mg, K, Ca, P, Mn, Fe, Cu, Zn, As, Hg, Tl and Pb in this solution was performed using a triple quadrupole inductively coupled plasma mass spectrometer (ICPMS, Agilent 8900, Santa Clara, California, USA) at the ICPMS labs, Dept. of Geology and Geophysics, University of Utah (https://earth.utah.edu/research_facilities/earth-core-facility/icp-ms.php). An external calibration curve was prepared from 1,000 mg/L standard (Inorganic Ventures, Christiansburg, VA, USA) with maximum concentrations of 10, 5, 2.5, 12, 0.2, 0.5, 0.04, 0.2, 0.04, 0.002 and 0.02 10 mg/mL for Na, Mg, K, Ca, Mn, Fe, Cu, Zn, As, Tl and Pb respectively. A second calibration curve was prepared for P and Hg, with maximum concentrations of 10 and 0.003 mg/mL respectively. Diluted samples, calibration solutions and blanks, were added 10 ng/mL in as internal standard and run in the ICPMS using a dual pass quartz spray chamber; PTFE nebulizer and dual-syringe introduction system (Teledyne, AVX 71000), platinum cones and sapphire injector in a platinum-shielded quartz torch. Oxygen mode was used in the reaction cell P determination, with masses 31>47. All other elements were determined using He mode at masses, 23, 24, 39, 44, 55, 56, 63, 66, 75, 202, 205 and 208 for Na, Mg, K, Ca, Mn, Fe, Cu, Zn, As, Hg, Tl and Pb respectively. Limit of determinations (LoD) for each element were calculated as three times the standard deviation of the blanks, multiplied by the total dilution factor used for samples (~4,000). Standard reference material 1643f (Trace Elements in Water, National Institute of Standards and Technology, Gaithersburg, MD, US) was run with calibration curve and samples (one standard every 10 samples) at a dilution of 1:20, reproducing certified values for Na, Mg, K, Ca, Mn, Fe, Cu, Zn, As, Tl and Pb within 5%. An in-house P standard solution prepared gravimetrically from high-purity $KH_2PO_4$ (Suprapur, Millipore-Sigma) with a concentration of 4 mg/mL, was run with P calibration curve and samples, reproducing the calculated value within 5%. The ICPMS instrument was located in a filtered air positive pressure lab and sample handling and dilutions were performed in laminar flow benches and using calibrated pipettors (Eppendorf Reference, Hamburg, Germany). All chemicals used were trace metal grade quality. Solution and dilution factors were used to calculate de concentration reported for each extracted element, in units of milligrams per kg of sediment.

Total Organic Carbon and Inorganic Carbon analyses

For carbon analyses, both methods used a TOC-L Carbon Analyzer connected to SSM-5000A furnace (Shimadzu, Kyoto, Japan). For total carbon (TC): 20-50 mg of sediment were measured in a ceramic crucible and covered with ceramic wool before being placed in the furnace at 900 ºC in an oxygenated atmosphere. For inorganic carbon (IC): 20-50mg of sediment were measured in a ceramic crucible and 0.5 mL of dilute phosphoric acid was added before being placed into the furnace at 250 ºC. Total organic carbon (TOC) is calculated from these two values, where TOC = TC - IC. Calibration curves were run before analyses using high purity $C_{12}H_{22}O_{11}$ (sucrose) and $NaHCO_3$ for TC and IC respectively.

Bacteria identification

To determine what bacterial communities are present in and on these worms, we performed a metagenomic study of the bacterial communities associated with individual nematodes collected from site 4, Fremont Island within the GSL. To identify and analyze bacterial associates, we amplified the V4 region of the 16S rRNA gene using the region-specific primer pair 515F and 806R, which included sequencer adapter sequences used in the Illumina MiSeq flow cell (*57*, *58*). We extracted bacterial DNA in 16 single worms prepared in 10 µL of single worm lysis buffer (pH 8.3: 121 mg Tris, 380 mg KCl, 51 mg $MgCl_2*6H_2O$, 460 µL Nonidet P-40, 460 µL Tween-20 per 100 mL $ddH_2O$) and incubated in the thermocycler (2 h at 65°C, 10 min at 95°C, and 4°C thereafter). We did not add Proteinase-K to the single worm lysate solution, as commonly done, since we found that DNA from worms lysed with Proteinase-K did not amplify well (Fig. S8). As negative sequencing controls, we used 8 samples from picks dipped in site-specific water samples but not transporting a worm. These negative controls did not show up in the gel, indicating there was not enough environmental DNA to amplify, arguing that sequenced microbes were specific to their worm hosts (Fig. S6). Each 60 µl PCR mixture (20-µl in triplicate) contained 6 µl TAQ buffer (NEB), 3 µl 5µM forward primer, 3 µl 5µM reverse primer, 1.2 µl 10 mM dNTP mix, 0.48 µl Taq polymerase (NEB), 5 µl of template DNA, and 41.32 µl sterile DNA-free PCR water. Multiplexing occurred at this step with six forward primers and one reverse primer. After making the master mix for each sample, we split it into three 20 µl reactions that ran in parallel. The PCR conditions were 94°C for 2 min, with 29 cycles at 94°C for 30 s, 50°C for 30 s, and 72°C for 60 s, with a final extension of 2 min at 72°C. We ran 10 µl of each sample on a gel to ensure 16S was amplified from our potentially low abundance template. After confirming the presence of bands after the first PCR, we took 1 µl per sample on to a second PCR to add the indexes. Each 25 µl PCR mixture contained 5 µl Q5 buffer, 0.125 µl 100µM Truseq forward primer, 1.25 µl 5µM barcoded reverse primer, 0.5 µl 10mM dNTP mix, 0.25 µl Q5 polymerase, 1 µl of DNA from the first PCR, and 16.875 µl sterile DNA-free PCR water. The PCR conditions were 94°C for 2 min, with 6 cycles at 94°C for 20 s, 60°C for 30 s, and 72°C for 30 min, with a final extension of 2 min at 72°C. We ran 5 µl of each sample on a gel to confirm that the barcodes were added and ensure each amplicon was ~440bp long. We next performed SPRI bead cleanup with 0.8x ratio to remove primer dimers and ran 5 µl of a 25 µl elute on a gel. We quantified dsDNA concentrations using a fluorometer (Qubit Broad Range sensitivity, Invitrogen) and determined average library sizes using a Tape Station and screen tape (Agilent Tech HS D100 Screen Tape). After quantification, the pool was diluted to 4nM (confirmed via Qubit), denatured, and then further diluted to a final concentration of 9pM with a 10% PhiX spike for sequencing. 16s rRNA gene amplicons were sequenced using multiplexed, paired-end 2x300 bp Illumina MiSeq sequencing. Cluster density was 486 $K/mm^2$; clusters passing filter was 98.7%; %>Q3 was 93.1%; and estimated yield was 7319.3 MB with $6.63 \times 10^6$ bacterial sequences.

Post-sequence processing was performed with DADA2 (*59*, *60*). The first 20 base pairs were truncated, as well as lower quality reads at the first instance of a Q score less than or equal to 2. Remaining forward reads were trimmed to 290 base pairs and reverse reads were trimmed to 250 base pairs. These high-quality sequences were dereplicated and denoised, then paired-end reads were merged. Next, we filtered chimeric reads from our sequence table and assigned taxonomy using the Silva database for classification of 16S sequences (*61*). Taxa present in less than 1% relative abundance were removed from subsequent analysis. After filtering, our datasets still generated $6.04 \times 10^6$ bacterial sequences from 15 individual nematodes.

Nematode identification

The isolated nematodes were further identified by morphology and molecular signatures. Morphological observations were made on live specimens, anesthetized using 20 mM sodium azide, on 2% agarose slides under a Zeiss Axiolab 5 microscope fitted with an Axiocam 208 color camera. For molecular analysis, individual worm lysate was prepared in single worm lysis solution (0.2 µL 10mg/mL Proteinase-K + 9.8 µL single worm lysis buffer pH 8.3: 121 mg Tris, 380 mg KCl, 51 mg $MgCl_2$*$6H_2O$, 460 µL Nonidet P-40, 460 µL Tween-20 per 100 mL $ddH_2O$). To extract DNA, we placed individual worms in 10 µL of single worm lysis solution and into the thermocycler (2 h at 65°C, 10 min at 95°C, and 4°C thereafter). Two of 11 attempted universal or nematode-specific primer pairs were successful in amplifying DNA of the expected size, both of which targeted the18S ribosomal small subunit (SSU rDNA) (Table S3). The gene fragments were amplified (*15*, *20*) and sequenced using standard PCR amplification conducted in 25 µl volumes with an initial denaturation step of 30 seconds at 95°C; 30 cycles of a 30 s denaturation at 95°C, 30 s annealing at 53°C, and 45 s extension at 68°C; and a final extension of 5 min at 68°C. We loaded 10 µl of each PCR product onto a 1% agarose gel to determine the size of the amplified product. Samples with visible bands of the expected size and purity were selected for sanger sequencing at the University of Utah DNA Sequencing Core Facility.

From aligned SSU sequences, we constructed Maximum likelihood phylogenetic trees of isolated nematodes (Fig. 4, Fig. S8). Sequence alignment was performed using MAFFT version 7.490 (*62*) and BMGE version 1.12 (*63*) with entropy score cut-off set to 0.6, gap-rate cut-off set to 0.3, and minimum block size set to 2. Analysis was performed in IQ-TREE multicore version 2.1.3, using a General Time Reversible (GTR) model for nucleotide substitutions (*64*). Nodal support for maximum likelihood analysis was performed for species identification with 1,000 bootstrap replicates. If the isolated nematode exhibited >97% identity to its best SSU BLAST hit (NCBI), that individual's tip got labeled with the relevant taxonomic information. To simplify the tree, clades consisting of multiple individuals of a single species were collapsed into a single branch and labelled with sample sizes and nodes with support values below 70 within a single family were collapsed into polytomies. Thus, the maximally inclusive subtree was selected where each taxon was represented by no more than one sequence or, in cases where more than one sequence was present for any taxon, all or most sequences from that taxon formed a clade or were part of the same polytomy, with only 4 exceptions to this rule that are indicated with dashed lines (Fig. 4).

Survival assay

We tested the viability of two common laboratory model nematodes in GSL water for five minutes. We placed *Caenorhabditis elegans* (N=59) and *Pristionchus pacificus* (N=64) adults into GSL water, assessed viability within five minutes of being placed and every 24 hours for four days, and transferred each individual back onto standard NGM plates. Similarly, we placed GSL nematodes (N=36) onto standard NGM plates and assessed viability within five minutes and at 24 hours later. Plates were incubated at 20°C and surviving animals were counted by their physiology and pick touch-provoked movement. Specifically, animals were counted as dead if they met at least two of the following criteria: lack of spontaneous or touch-evoked swimming, gray discoloration, tissue degradation, body wall rupturing, and stiff, especially straight posture (*20*).

Statistics

Nematode abundance across samples was log10(n+1) transformed. Univariate ANCOVAs were performed using site as the categorical variable and salinity as a covariate. An ANOVA was performed using season within 2021 (spring, summer, fall) as the categorical variable. Mann-Whitney Wilcoxon tests were performed on the subset of samples collected from GSL using microbialite (vs. non-microbialite) and year (2021 vs. 2022) as a categorical variable. Univariate ANCOVAs were performed on the subset of samples collected from GSL using microbialite status (microbialite vs. non-microbialite) as the categorical variable and pH, soil moisture content, or oxidation reduction potential as a covariate.

For mass spectrometry data of the chemical environment in and around GSL, we performed Mann-Whitney Wilcoxon tests on the subset of samples collected from GSL using microbialite (vs. non-microbialite) as a categorical variable to see if there was a microbialite effect on the chemical environment. We also performed Mann-Whitney Wilcoxon tests on data from all sites to see if there was an effect of salty vs. freshwater sites on each component of the chemical environment. Univariate ANCOVAs were performed using season as the categorical variable and salinity as a covariate on each component of the chemical environment. ANOVAs were performed to see if chemical environment influenced nematode abundance. All statistical tests were carried out in the R statistical environment (version 4.1.2, R Development Core Team 2019, http://www.r-project.org) in RStudio (version 1.4.869, RStudio Team 2019).

### SUPPLEMENTARY FIGURES & CAPTIONS

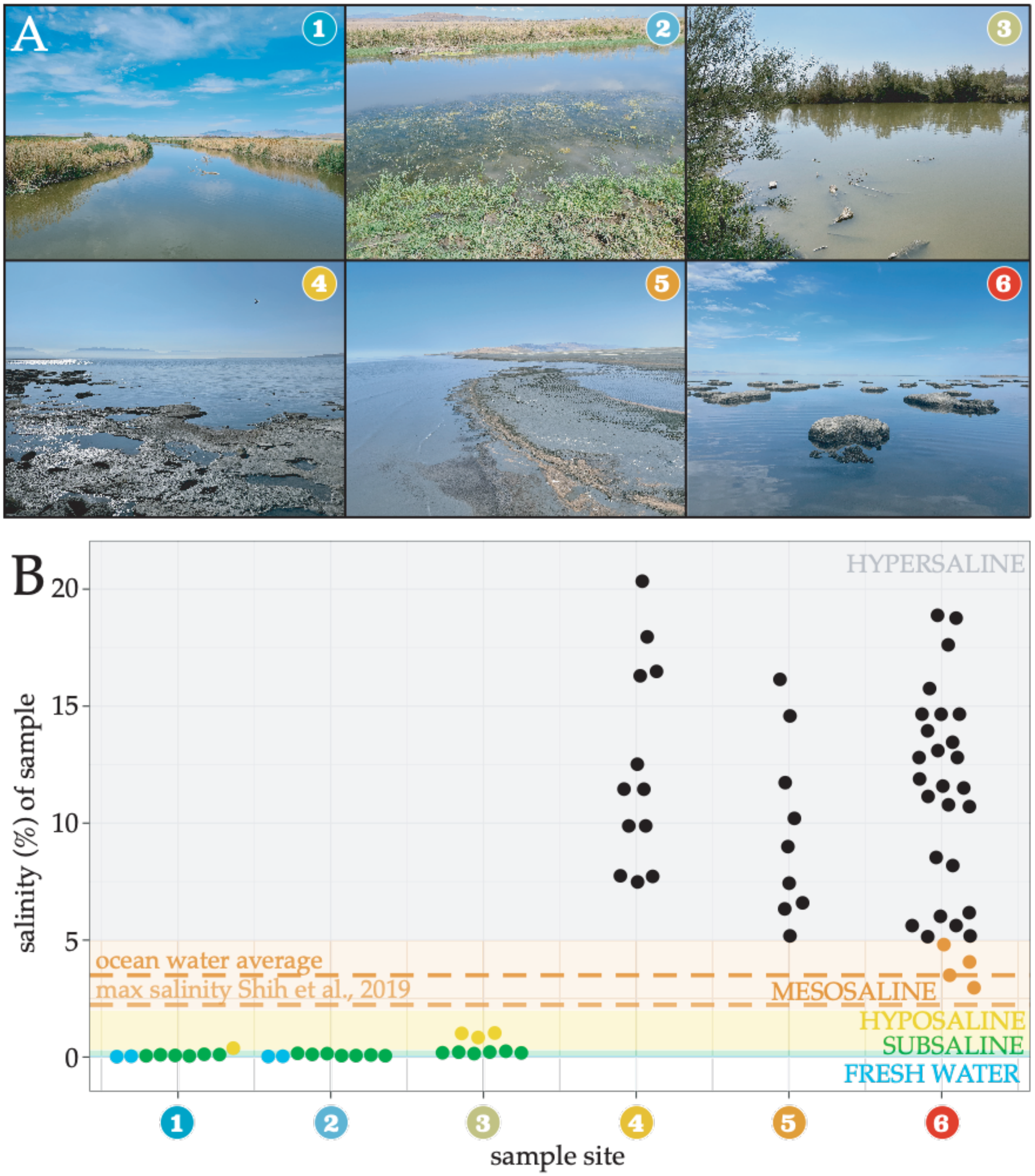

**Fig. S1. The Great Salt Lake is hypersaline. (A)** Photographs of the six sampling sites around (1-3) and within (4-6) the Great Salt Lake. Samples were collected in spring, summer, and fall of 2021 and the summer of 2022. **(B)** Salinity values of lake sediment samples from each site in percent. Data points represent individual samples and are spread horizontally to show stacked points.

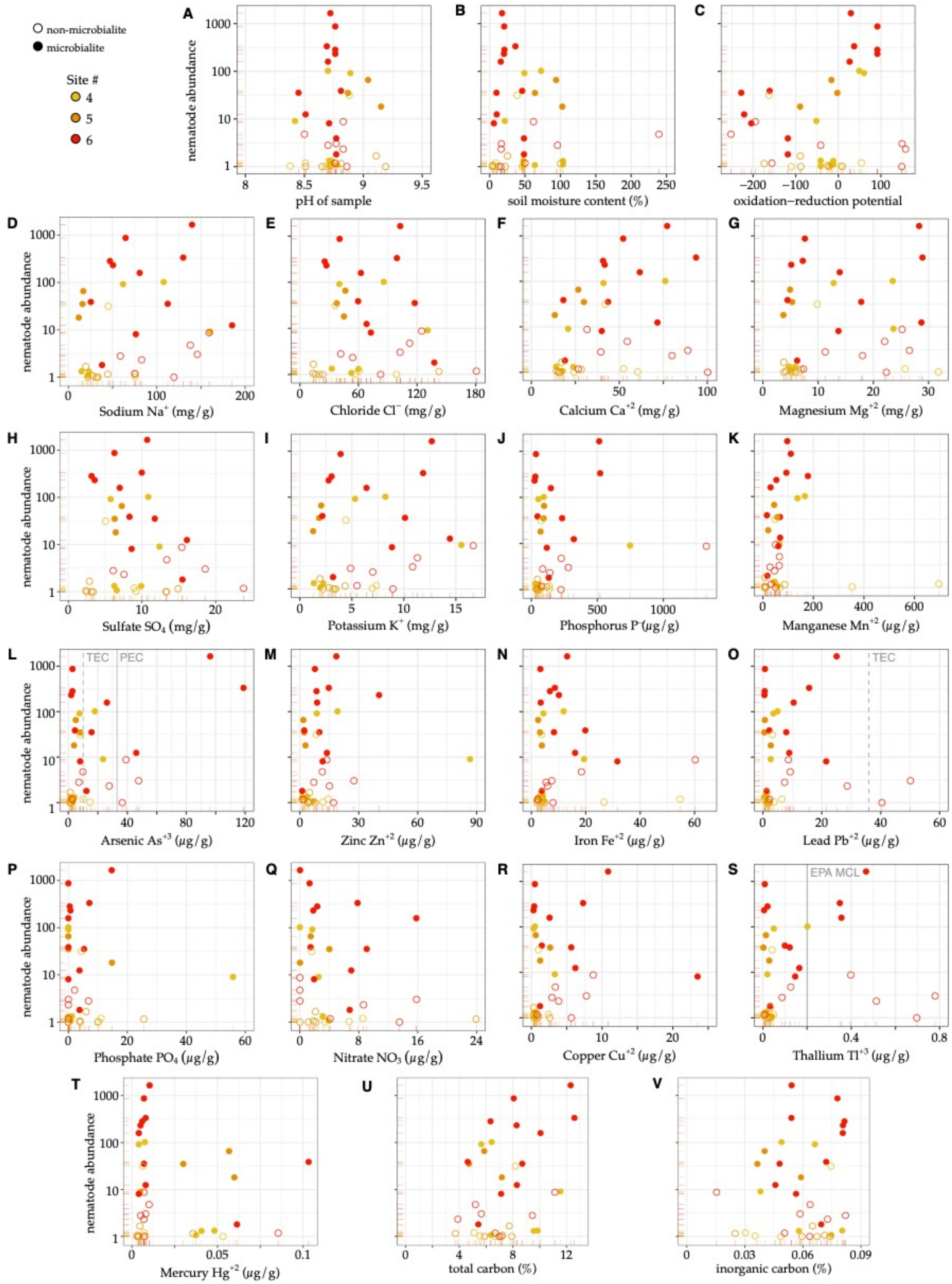

**Fig. S2. Chemical composition of soil samples from within the Great Salt Lake.** Nematode abundance by **(A)** pH, **(B)** soil moisture content, **(C)** oxidation-reduction potential, **(D)** sodium,

**(E)** chloride, **(F)** calcium, **(G)** magnesium, **(H)** sulfate, **(I)** potassium, **(J)** phosphorus, **(K)** manganese, **(L)** arsenic, **(M)** zinc, **(N)** iron, **(O)** lead, **(P)** phosphate, **(Q)** nitrate, **(R)** copper, **(S)** thallium, **(T)** mercury, **(U)** total carbon, and **(V)** inorganic carbon content of each sample. Abundance is measured in number of nematodes per 100g of dry lake sediment on a log scale. Data points each represent a soil sample from which nematodes were extracted, colored by the site from which the sample was collected. Filled points represent microbialite samples and unfilled points represent non-microbialite samples. Total carbon is the sum of carbon species in the sample, including both inorganic and organic carbon.

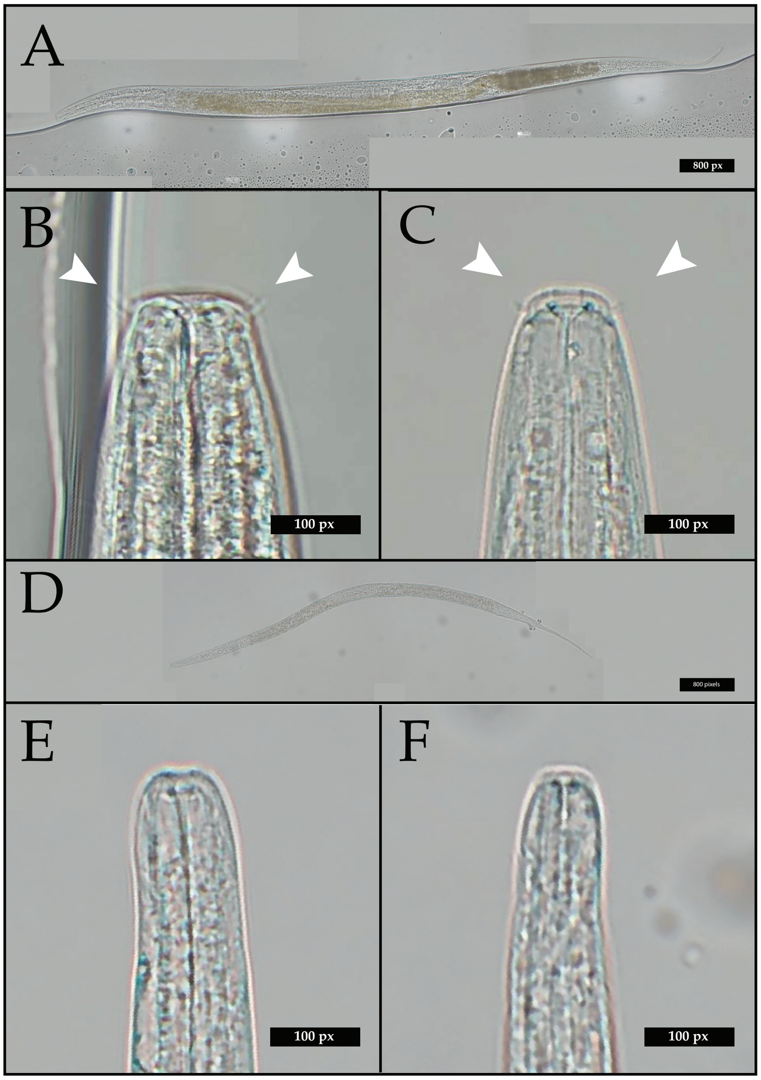

**Fig. S3. Great Salt Lake nematodes are likely bacteriovores.** **(A)** Body and **(B–C)** mouths of Monhysteridae worms isolated from fresh water as compared to **(D)** body and **(E–F)** mouths of Monhysteridae worms isolated from hypersaline sites within the Great Salt Lake. Black bars show 800 pixels in body shots and 100 pixels in head shots. White arrows highlight conspicuous cephalic setae shared amongst freshwater Monhysteridae.

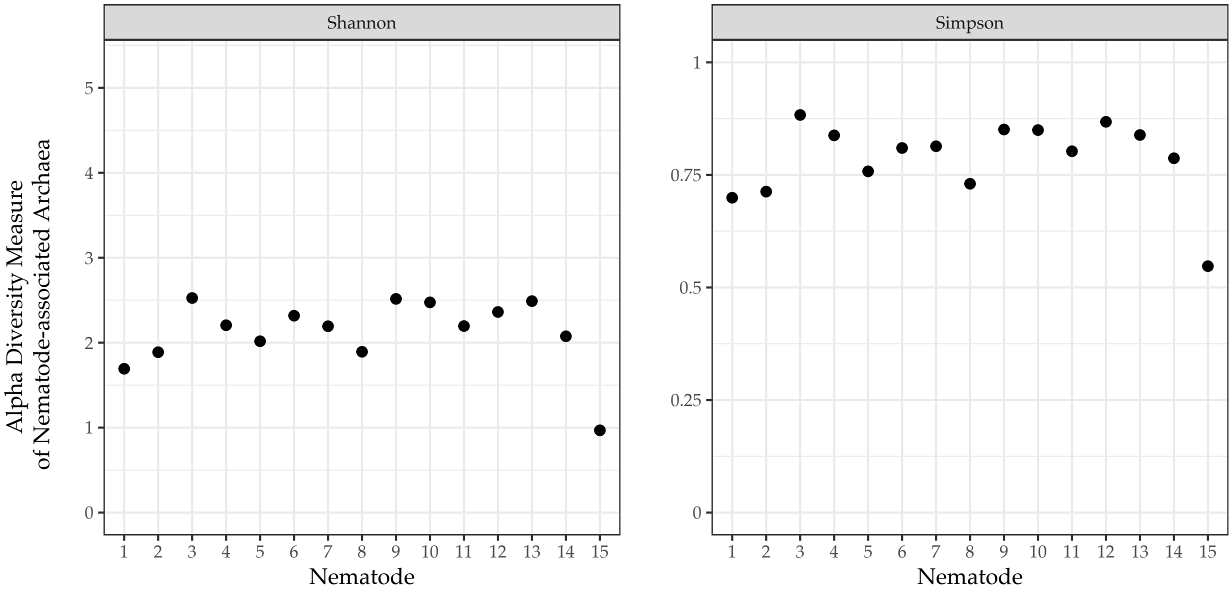

**Fig. S4. Alpha diversity indices of nematode-associated archaea.** Each dot represents an individual nematode from which 16S data were analyzed. All 15 nematodes were collected from Fremont Island (site 4) and taxonomic bins are grouped by phylum.

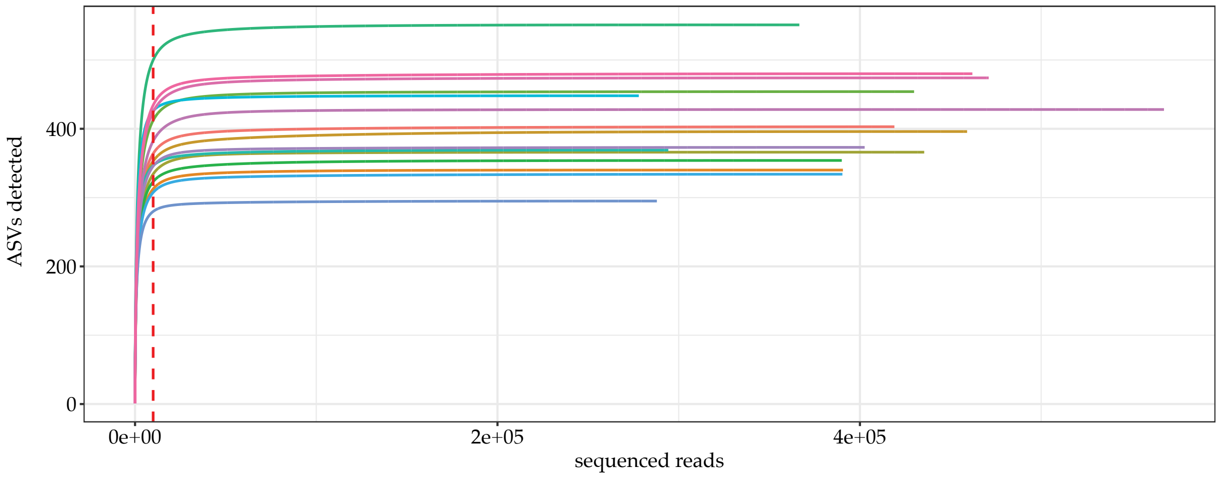

**Fig. S5. Our analysis of nematode-associated microbiota captures much of the diversity in our samples.** Rarefaction curves show the number of Amplicon Sequence Variants (ASVs) detected as a function of sequencing depth. They are used to estimate whether all the diversity of the true community was captured. Each colored line represents a different nematode sample and shows how many ASVs are found in a random subset of different numbers of reads. For our samples, few new ASVs are observed after about 10,000 reads (vertical red dashed line), suggesting that the samples had sufficient reads to capture most of the diversity or reach a point of diminishing returns. All 15 nematodes were collected from Fremont Island (site 4) and taxonomic bins are grouped by phylum.

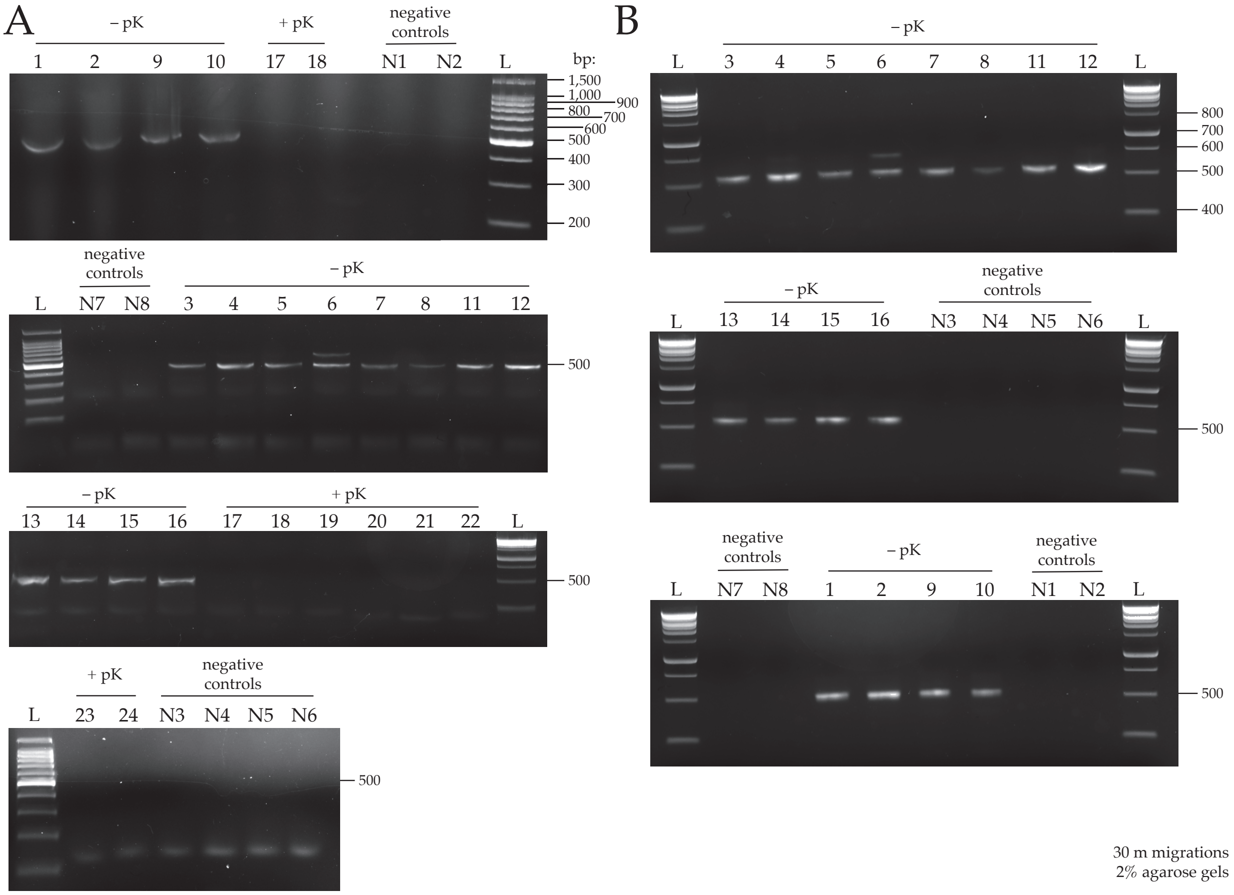

**Fig. S6. 16s rRNA amplicons of the V4 region in 24 sample individual worms and 8 negative controls.** Amplicons are shown after the **(A)** first and **(B)** second PCR reactions, visualized after electrophoretic separation. Samples 1-16, for which DNA was extracted without the addition of Proteinase-K (– pK), showed a band at a reasonable amplicon size ~500-bp, while samples 17-24, for which DNA was extracted with the addition of Proteinase-K (+ pK), did not show a band after the first PCR reaction. None of the negative controls (N1-N8) showed any bands, indicating insufficient DNA to sequence. As a result, we proceeded in the analysis with samples extracted without Proteinase-K. All runs were 30 min migrations on 2% agarose gels. L: DNA size marker, either 100 bp ladder or 1Kb Plus ladder.

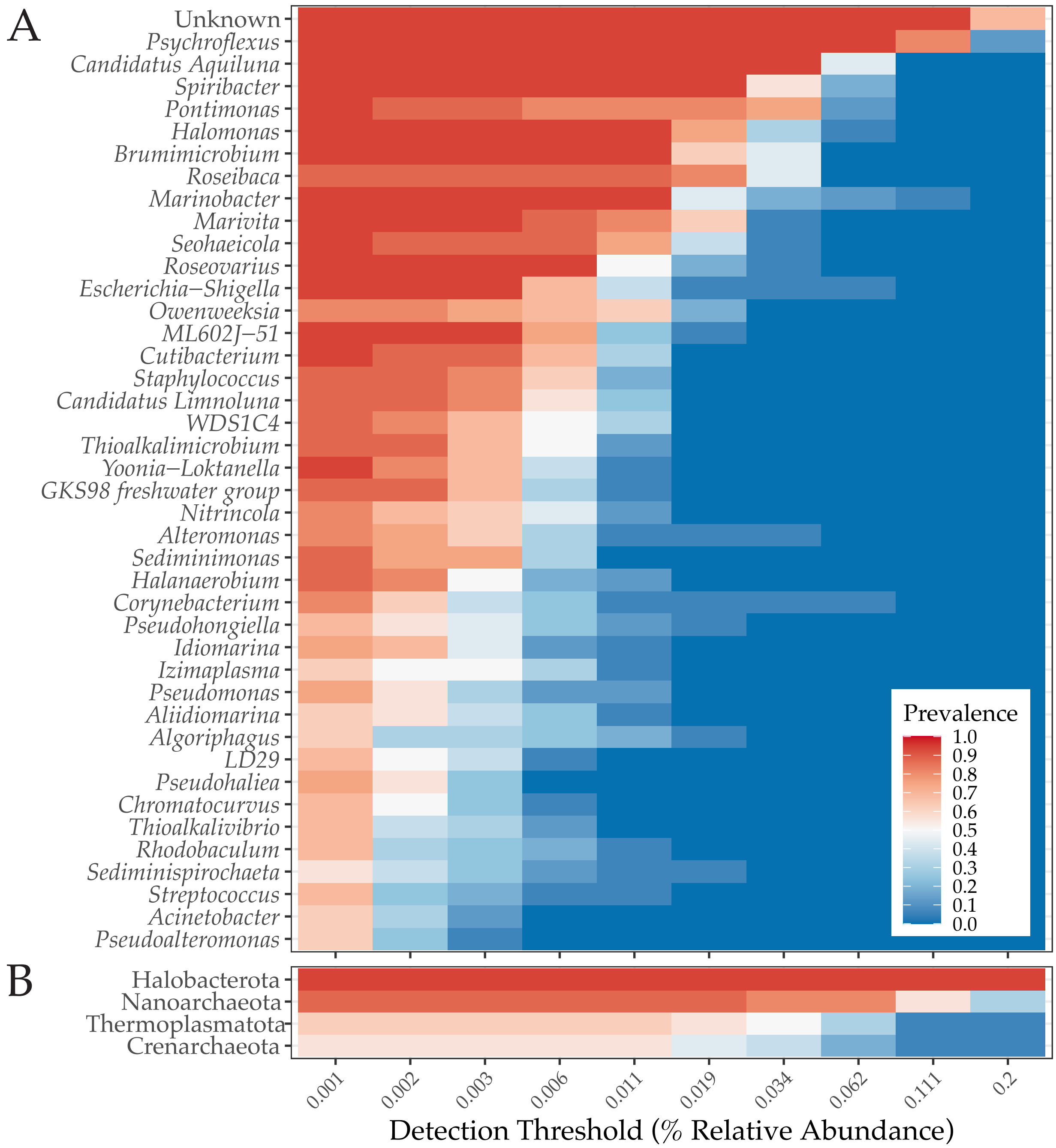

**Fig. S7. Core microbiota.** The set of **(A)** bacterial genera and **(B)** archaeal phyla shared by almost all individual nematodes. All taxa are transformed to compositional or relative abundances with a minimum prevalence of 95%.

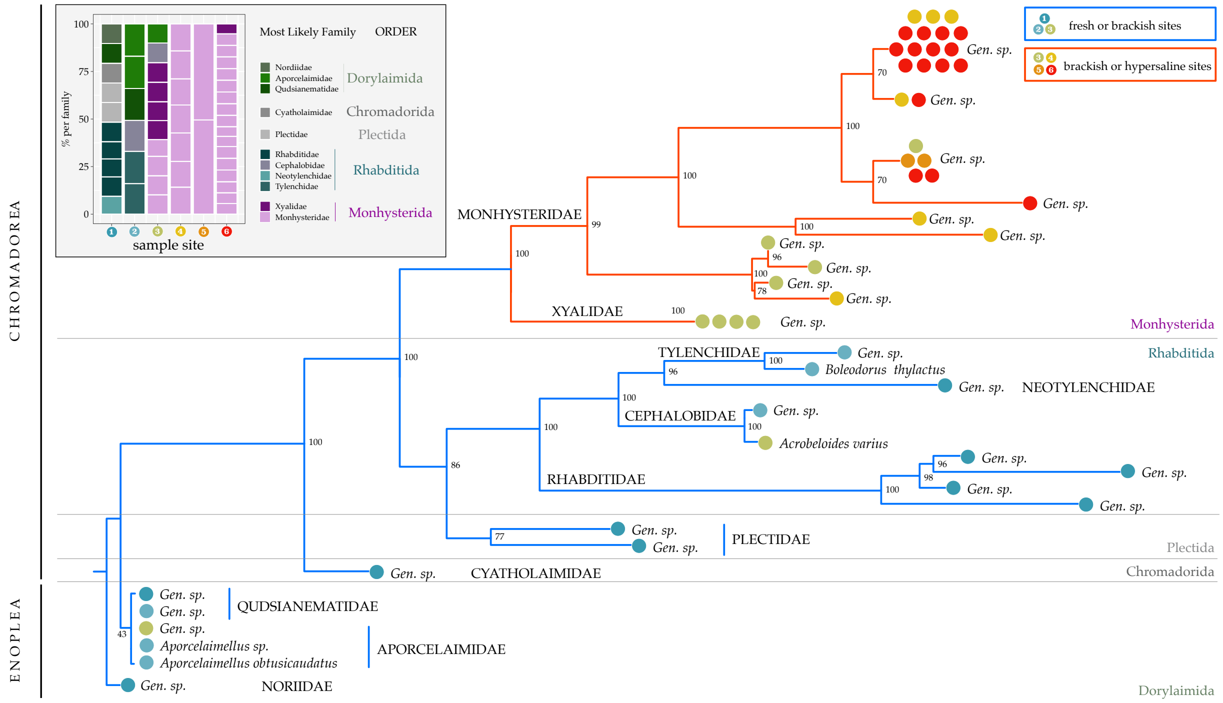

**Fig. S8. Phylogenetic tree of Great Salt Lake nematodes using a second primer set.** Nematodes were isolated and underwent single-worm lysis and PCR with a second set of SSU primers. PCR samples with single bands on a 1% agarose gel were sent for Sanger Sequencing at the University of Utah DNA Sequencing Core Facility. Aligned reads (MAFFT) were used as input for a Maximum Likelihood phylogenetic tree using a GTR model for nucleotide substitutions. Nodal support (shown in colored numbers) for analysis was performed for species identification with 1,000 bootstrap replicates. If a nematode exhibited >97% identity to its best SSU BLAST hit (NCBI), we labeled its tip with the relevant taxonomic information. Clades consisting of multiple individuals of a single species were collapsed into a single branch and labelled with sample sizes. If percent identity exceeded 97%, that individual's tip was labelled by the scientific name of the closest relative. To simplify the tree, clades that were home to individuals within a single species were simply labelled with that species' name and nodes with support values below 0.70 within a single family were collapsed into polytomies. **(Inset)** Nematode family composition by site. Each box is an individual nematode, colored by the most likely family and order.

### SUPPLEMENTARY TABLES & CAPTIONS

**Table S1. Global nematode abundances per 100g dry soil.**

| Global nematode abundances per biome per 100g dry soil | | | | |
|---|---|---|---|---|
| Biome | Mean | Median | Samples | Sites |
| Tundra | 5196.31625 | 2329.405 | 148 | 22 |
| Boreal Forests | 3440.217856 | 2159.636364 | 669 | 201 |
| Temperate Broadleaf Forests | 3401.142253 | 2076.5 | 2109 | 919 |
| Tropical Coniferous Forests | 999.75 | 999.75 | 8 | 1 |
| Temperate Conifer Forests | 2539.721373 | 906 | 173 | 93 |
| Montane Grasslands | 3645.784738 | 628 | 116 | 39 |
| Tropical Grasslands | 640.8987288 | 567.25 | 271 | 60 |
| Tropical Moist Forests | 1032.573342 | 475.5 | 969 | 197 |
| Temperate Grasslands | 842.9087684 | 445.7666667 | 627 | 92 |
| Mediterranean Forests | 677.1121939 | 430.3391667 | 703 | 151 |
| Tropical Dry Forests | 281.55 | 281.55 | 11 | 2 |
| Flooded Grasslands & Savannas | 183.2857143 | 183.2857143 | 7 | 1 |
| Antarctica | 1274.514053 | 96.4 | 503 | 39 |
| Deserts | 224.9074426 | 81.28571429 | 346 | 75 |
| Owens Lake | 50.44521 | 28.52873 | 8 | 5 |
| Weber River Input | 39.115 | 19.949 | 27 | 3 |
| Great Salt Lake | 113.706 | 7.088 | 35 | 3 |
| Mono Lake | 24.57509447 | 6.189043843 | 28 | 3 |
| Global biome nematode abundance values are from van den Hoogen et al. 2019 | | | | |
| Owens Lake, Weber River Input, and Great Salt Lake are our comparisons, reported herein | | | | |
| Mono Lake nematode abundance is from Shih et al. 2019 | | | | |

**Table S2. Statistical analysis of physiochemical profiles of soil samples.** Specific statistical tests used are detailed in Methods. Values shown are P values for the fixed effect of site, salinity, season, and microbialite state on each aspect of the chemical environment, as well as the fixed effect of each aspect of the chemical environment on nematode abundance. Microbialite analysis is on GSL sites only (sites 4-6), while all other analyses are on all sites (1-6).

|  | Fixed Effect of ------ on Ion or Compound: (P) |  |  |  |  | Fixed Effect of Ion or Compound on ------: (F, P) |  |
|---|---|---|---|---|---|---|---|
| | Great Salt Lake or Freshwater | Salinity (% of Sample) | Season (Spring, Summer, or Fall) | Salinity:Season Interaction | Microbialite | Nematode Abundance | |
| Sodium ($Na^+$) | **< 2.2e-16** | **1.238e-07 ***** | **0.040093 *** | **0.007943 **** | 0.7291 | 0.0545 | 0.8162 |
| Chloride | **< 2.2e-16** | **9.35e-09 ***** | 0.1035 | 0.2505 | 0.7291 | 1.7005 | 0.197 |
| Potassium | **2.16E-15** | **4.169e-11 ***** | **0.003062 **** | **0.007549 **** | 0.4065 | 0.0055 | 0.9413 |
| Calcium | 0.4492 | 0.05826 . | **5.194e-10 ***** | 0.81329 | 0.9161 | 2.8978 | 0.09371 . |
| Phosphorus | 0.198 | **0.001308 **** | **0.003396 **** | **0.008564 **** | 0.7519 | 0.6246 | 0.4323 |
| Magnesium | **4.61E-16** | **3.52e-12 ***** | **0.001151 **** | **0.024416 *** | 0.7066 | 5e-04 | 0.9827 |
| Manganese | **3.36E-08** | **0.0007938 ***** | **0.0013346 **** | 0.1169753 | 0.1582 | 0.2371 | 0.628 |
| Zinc | **6.04E-06** | **0.0001314 ***** | **2.770e-08 ***** | **2.399e-05 ***** | 0.964 | 2.954 | 0.09066 . |
| Iron | **6.15E-05** | **0.007014 **** | **2.705e-05 ***** | 0.536329 | 0.5567 | 0.1909 | 0.6637 |
| Total Carbon | **1.42E-11** | **< 2e-16 ***** | 0.07285 . | 0.5413 | 0.1408 | 0.0823 | 0.7752 |
| Inorganic Carbon | **< 2.2e-16** | **4.953e-11 ***** | 0.1947 | 0.8851 | 0.598 | 0.6386 | 0.4273 |
| Nitrate | **2.89E-01** | 0.570506 | **0.003691 **** | **0.002143 **** | 0.8908 | 1.7376 | 0.1923 |
| Sulfate | **< 2.2e-16** | **6.535e-11 ***** | 0.07856 . | 0.16076 | 1 | 3.4666 | 0.06736 . |
| Phosphate | **2.44E-09** | **0.0117333 *** | 0.5304202 | **0.0002221 ***** | 0.2496 | 0.2618 | 0.6107 |
| Mercury | 0.06357 | **0.005662 **** | **< 2.2e-16 ***** | **8.776e-07 ***** | 0.1639 | 0.6297 | 0.4305 |
| Copper | 0.265 | 0.6345627 | **0.0001118 ***** | 0.7206175 | 0.8909 | 0.5152 | 0.4756 |
| Arsenic | **2.44E-09** | **2.187e-14 ***** | **0.004924 **** | 0.687258 | 0.3413 | **4.6629** | **0.0347 *** |
| Lead | **1.52E-09** | **5.666e-10 ***** | **4.470e-05 ***** | 0.2943 | 0.2439 | 0.0058 | 0.9396 |
| Thallium | 0.746 | 0.05997 . | **8.086e-06 ***** | 0.2271 | 0.6623 | 0.0875 | 0.7683 |
| | Signif. codes: 0 ‘***’ 0.001 ‘**’ 0.01 ‘*’ 0.05 ‘.’ 0.1 ‘ ’ 1 | | | | | | |

**Table S3. Oligonucleotides for sequencing nematodes.** Of the 11 universal primer sets tested, two sets worked, which are bolded and marked with an asterisk.

| Oligonucleotides for sequencing nematodes | | |
|---|---|---|
| Gene marker/region | Forward or Reverse | Sequence (5' to 3' end) |
| ***SSU (18S) **** | ***F*** | ***AAAGATTAAGCCATGCATG*** |
| | ***R*** | ***CATTCTTGGCAAATGCTTTCG*** |
| ***SSU (18S) **** | ***F*** | ***CGCGAATRGCTCATTACAACAGC*** |
| | ***R*** | ***GGGCGGTATCTGATCGCC*** |
| SSU (18S) | F | GCTTGTCTCAAAGATTAAGCC |
| | R | TGATCCWMCRGCAGGTTCAC |
| LSU (28S) | F | ACAAGTACCGTGAGGGAAAGTTG |
| | R | TCGGAAGGAACCAGCTACTA |
| LSU (28S) | F | TTCGACCCGTCTTGAAACACG |
| | R | TCCTCGGAAGGAACCAGCTACTA |
| ITS1 | F | TTGATTACGTCCCTGCCCTTT |
| | R | TTTCACTCGCCGTTACTAAGG |
| ITS | F | CTTTGTACACACCGCCCGTCGCT |
| | R | TTTCACTCGCCGTTACTAAGGGAATC |
| IGS | F | TTAACTTGCCAGATCGGACG |
| | R | TCTAATGAGGGAACCAGCTACTA |
| cox1 | F | TGTCTTTACCWGTTTTRGCTGG |
| | R | CCGAAAGCAGGYAAAATHARAA |
| cox1 | F | GATTTTTTGGKCATCCWGARG |
| | R | GTACATAATGAAAATGTGCCAC |
| cox1 | F | TGAGGCATTCTATTATATATATGTCGG |
| | R | AGCAACAACATAGTAAGTATCGTG |
| * Primers that worked | | |

### SUPPLEMENTARY MOVIE CAPTION

**Movie S1.** Representative nematode extracted from the south arm of the Great Salt Lake on March 3, 2021.
